# Effects of Cholesterol Modulation on Cisplatin-Induced Hearing Loss

**DOI:** 10.1101/2025.09.23.678043

**Authors:** John Lee, Marcello Peppi, Megan Guidry, Katharine Fernandez, Abu Chowdhury, Lizhen Wang, Lisa L. Cunningham

## Abstract

Cisplatin is a widely used and effective anticancer drug. However, it causes permanent sensorineural hearing loss in over 50% of treated patients. There are no FDA-approved therapies to prevent cisplatin-induced hearing loss (CIHL) in adults, highlighting a critical unmet clinical need. Previous studies suggest that statins, commonly prescribed cholesterol-lowering drugs, are associated with reduced incidence and severity of cisplatin-induced hearing loss. Statins are primarily used to lower cholesterol, but they also exert several pleiotropic effects, making the mechanism(s) underlying this protection unclear. Here we examine whether reduced plasma cholesterol confers protection against CIHL independent of statin treatment. We utilized mice lacking serine protease proprotein convertase subtilisin/kexin type 9 (*Pcsk9* knockout (KO) mice) as a genetic model of reduced plasma cholesterol. We find that *Pcsk9 KO* mice are protected against cisplatin-induced hearing loss, as reflected by significantly lower ABR and DPOAE threshold shifts relative to wild type (WT) mice following treatment. Histological analyses confirmed preservation of cochlear outer hair cells (OHCs) in *Pcsk9 KO* mice treated with cisplatin, whereas WT mice showed significant OHC loss in the high-frequency cochlear regions. Finally, hearing loss positively correlated with baseline plasma cholesterol levels. Together our data demonstrate that systemic cholesterol reduction provides significant protection against CIHL, and they suggest that the protective effect of statins against CIHL is mediated by cholesterol reduction.

## 1. INTRODUCTION

Cisplatin is a first-line chemotherapeutic agent in the treatment of various tumors (Boulikas & Vougiouka, 2004). Its widespread use stems from its demonstrated efficacy in improving disease-free and overall survival in cancer patients (Einhorn, 2002). However, cisplatin has several significant toxicities (Barabas et al., 2008; Fung et al., 2018). Among these, cisplatin-induced hearing loss (CIHL) stands out as a serious and irreversible adverse effect, affecting up to 50% of treated adults (Fernandez et al., 2021; Frisina et al., 2016; Rybak et al., 2019)Cisplatin induces degeneration of several cell types in the cochlea, including outer hair cells (OHCs), spiral ganglion neurons, and cells of the stria vascularis (Callejo et al., 2015). CIHL can impair communication, hinder social interactions, and negatively impact quality of life for cancer survivors (Dillard et al., 2022; Gurney et al., 2007; Moke et al., 2021). There are currently no FDA-approved therapies to prevent or mitigate cisplatin-induced ototoxicity in adults with cancer, underscoring a pressing clinical need.

Statins, widely prescribed cholesterol-lowering drugs, function primarily by suppressing the activity of HMG-CoA reductase, a key enzyme central to endogenous cholesterol biosynthesis, leading to reduced plasma levels of LDL-C , aka “bad cholesterol”, the lipoprotein responsible for transporting cholesterol in the blood (Istvan & Deisenhofer, 2001). In addition to cholesterol lowering, statins also have anti-inflammatory and antioxidant effects that may confer cellular protection and contribute to the overall cardiovascular protection provided by statins (Liao & Laufs, 2005; Oesterle et al., 2017). We previously showed that lovastatin reduces cisplatin-induced hearing loss (CIHL) in mice (Fernandez et al., 2020) and that statin use is associated with reduced incidence and severity of CIHL in adults with head and neck cancer (Fernandez et al., 2021). These studies suggest that cholesterol lowering and/or other pleiotropic effects of statins may play a protective role in CIHL. A randomized controlled clinical trial (ClinicalTrials.gov ID: NCT04915183) is underway to determine whether atorvastatin, the most prescribed statin in the US (Matyori et al., 2023), may be repurposed to reduce CIHL in adults with head and neck cancer.

The mechanisms underlying the observed protective effect of statins against CIHL are unknown. Thus, to begin to understand the mechanisms underlying statin-induced protection, we asked whether reduced cholesterol in the absence of statins is also protective. Proprotein convertase subtilisin/kexin type 9 (PCSK9), a serine protease encoded by the *Pcsk9* gene plays a pivotal role in lipid metabolism by accelerating the degradation of low-density lipoprotein (LDL) receptors on hepatocytes (Rashid et al., 2005; Roubtsova et al., 2022). These LDL receptors function by binding cholesterol-rich low-density lipoproteins and internalizing them via endocytosis. When PCSK9 binds to LDL receptors, it redirects them toward lysosomal degradation, thus reducing the pool of LDL receptors and elevating plasma LDL cholesterol levels (Abifadel et al., 2003; Allard et al., 2005). Therapeutic inhibition of PCSK9 can be achieved through monoclonal antibodies (alirocumab and evolocumab) and RNA interference (inclisiran). These drugs are FDA approved and are effective at reducing cholesterol in patients at high cardiovascular risk (Sabatine et al., 2017).

Here we utilized *Pcsk9* knockout mice, which exhibit reduced plasma cholesterol due to the absence of PCSK9, as a genetic model for examining the effects of cholesterol modulation on CIHL. We reasoned that if the protective effect of statins is mediated by statin-induced reduction in plasma cholesterol, then genetically reducing cholesterol should similarly protect against CIHL.

## 2. MATERIALS AND METHODS

### Animals

All mice were housed individually with free access to food and water in accordance with the NIH NIDCD/NINDS Animal Care and Use Committee-approved protocol (#1327).

*Pcsk9 KO* mice (B6;129S6-Pcsk9tm1Jdh/J; The Jackson Laboratory, Bar Harbor, Maine, USA) carry a deletion in the *Pcsk9* gene that includes a portion of the 3′ region of exon 2 and all of exons 3 and 4, (Horton 2005). C57BL/6 (B6) mice, the background strain of the *Pcsk9 KO* mice are known to develop early-onset age-related hearing loss due to a mutation in the Cdh23 gene termed *Cdh23^ahl^* (Kane et al., 2012). To eliminate the confounding influence of hearing loss caused by *Cdh23^ahl^*, we crossed *Pcsk9 KO* mice with *Cdh23^ahl^*-corrected B6.CAST-Cdh23Ahl+/Kjn mice (Strain #: 002756), a strain known to maintain normal auditory function well into advanced ages. Offspring heterozygous for the *Pcsk9 KO* allele and heterozygous for the wildtype *Cdh23^ahl+^* allele were intercrossed to generate Pcsk9 wildtype (WT) and KO littermates on a B6 background free from the early-onset hearing loss *Cdh23^ahl^* allele.

### Atorvastatin and cisplatin treatments

Adult CBA/CaJ mice from the Jackson Laboratory (Bar Harbor, Maine, USA) were randomly assigned to one of four treatment groups, maintaining balanced sexes across groups: (1) saline control, (2) cisplatin only, (3) cisplatin + 40 mg/kg atorvastatin, or (4) cisplatin + 60 mg/kg atorvastatin. Adult CBA/CaJ mice underwent cisplatin treatment using a previously described protocol (Fernandez et al., 2019): three cycles of once-daily intraperitoneal (IP) injections of cisplatin (3 mg/kg) for 4 days, each followed by a 10-day recovery period (Figure 1A), totaling 42 days. Atorvastatin calcium (1044516-100MG, Millipore Sigma, USA) was dissolved in water (vehicle) via oral gavage (20G, Roboz FN-7910), beginning 24 hrs prior to cisplatin administration and then once daily throughout the cisplatin protocol. Cisplatin-only treated mice were given vehicle via oral gavage at volumes equivalent to the higher dose of atorvastatin (60 mg/kg). All cisplatin-treated CBA/CaJ mice received daily supportive care including: 1) subcutaneous injections of 1 ml 0.9% NaCl in the morning and 1 ml of Normasol (Hospira Inc., San Clemente, CA, USA) in the afternoon, and 2) 0.3 ml of a high-calorie or high-protein oral nutritional supplement to maintain body weight. Supplement options included STAT® liquid high-calorie supplement (PRN Pharmacal, Pensacola, FL, USA).

**Figure 1.**
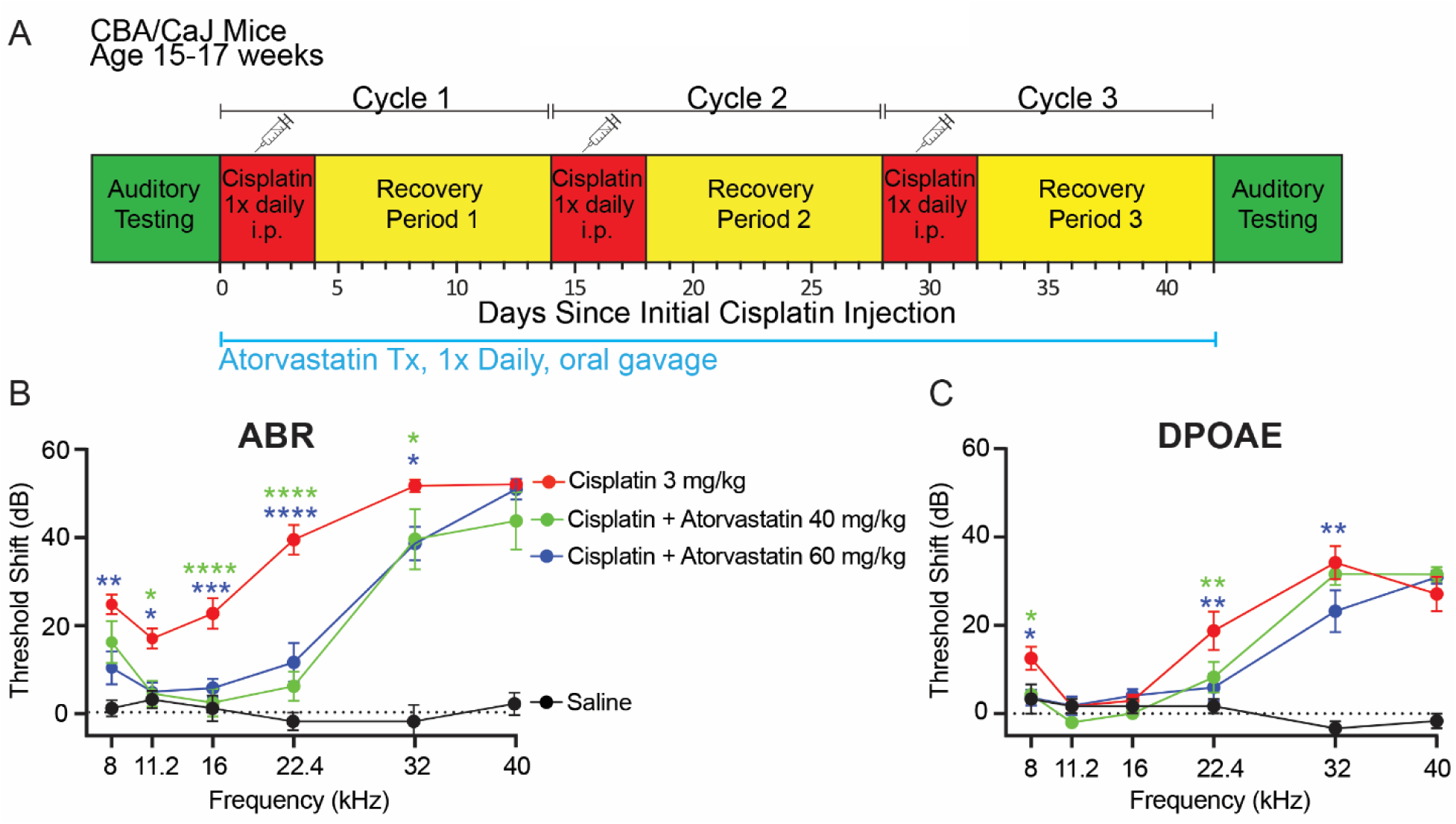
Atorvastatin reduces cisplatin-induced hearing loss (CIHL) and OHC dysfunction in mice. **(A)** A multi-cycle cisplatin administration protocol was used to characterize CIHL in mice with and without atorvastatin. **(B)** ABR threshold shifts demonstrate that cisplatin treatment resulted in significant hearing loss across frequencies (n = 29). Both doses of atorvastatin (60 mg/kg and 40 mg/kg, n = 10-12 for each group) resulted in significant protection against CIHL (p ≤ 0.05, two-way ANOVA). Saline-treated control mice (n = 9) had no hearing loss. **(C)** DPOAE threshold shifts demonstrate that cisplatin-treated mice had significant reduction in OHC function. Both doses of atorvastatin were partially protective against cisplatin-induced shifts in DPOAE thresholds (n = 11-13 for each group, p ≤ 0.05, two-way ANOVA). Data are expressed as mean ± SEM [Green asterisks: Cisplatin vs. Cisplatin + Atorvastatin 40 mg/kg; blue asterisks: Cisplatin vs. Cisplatin + Atorvastatin 60 mg/kg; as determined by two-way ANOVA, * p ≤ 0.05, ** p ≤ 0.01, *** p ≤ 0.001, **** p ≤ 0.0001].

Additional food pellet chow and DietGel® Recovery cups (ClearH2O, Portland, ME, USA) were also provided. Mice were monitored daily by investigators and veterinary staff for changes in overall health, and body conditioning scores (BCS) were recorded to assess muscular tone, body fat, coat maintenance, and energy level (Ullman-Cullere & Foltz, 1999).

As *Pcsk9 KO* mice are on a C57BL/6 genetic background—which our unpublished data indicate is slightly less susceptible to CIHL—we modified the cisplatin administration protocol by increasing the individual cisplatin dose from 3 mg/kg to 3.5 mg/kg. Forty-two adult Pcsk9 mice (21 males, 21 females) were randomly assigned to saline control (n = 15) or cisplatin-treated (n = 27) groups. Supportive care was provided as described for CBA/CaJ mice, with the following exception: instead of STAT® supplement, *Pcsk9 KO* mice received a custom blend of ZooPro High Protein Supplement and Ensure High Protein Nutrition Shake (Exotic Nutrition Pet Supply, Newport News, VA, USA), along with Ensure High Protein Milk Chocolate Nutrition Shake (Abbott Nutritional Products, Abbott Park, IL, USA) daily to support maintenance of body weight. All cisplatin-treated *Pcsk9 KO* mice also received daily subcutaneous injections of 1 ml 0.9% NaCl in the morning and 1 ml of Normasol in the afternoon, along with food pellet chow and DietGel® Recovery cups. Mice were monitored by investigators and veterinary staff for changes in overall health that may have resulted from cisplatin treatment. Body conditioning scores (BCS) were recorded daily and used to monitor the overall condition based on muscular tone, body fat content, coat maintenance, and overall energy level (Ullman-Culleré & Foltz 1999).

### *Pcsk9* Genotyping

Genomic DNA was extracted from mouse tail biopsies by overnight digestion in 20 mg/mL proteinase K (cat# 503-PK, Viagen, St. Louis, MO) in DirectPCR lysis reagent (cat# 102-T, Viagen, Los Angeles, CA), following the manufacturer’s instructions. PCR-based genotyping was performed using 1 µL of extracted DNA and primers designed to detect wild-type and mutant alleles of the *Pcsk9* locus. For genotyping the endogenous *Pcsk9* gene, standard PCR was carried out using the following primers: 5′- ATTGTTGGAGGGAGAAGTACAG-3′, 5′-GCGAGCATCAGCTCTTCATA-3′, and 5′-GATTGGGAAGACAATAGCAGGCATGC-3′. Thermal cycling conditions included an initial denaturation at 95°C for 3 minutes, followed by 29 cycles of denaturation at 95°C for 30 seconds, annealing at 61°C for 30 seconds, and extension at 72°C for 1 minute. PCR products were resolved on 2% E-Gel™ agarose gels (cat# G800802, Invitrogen, Waltham, MA, USA), where a 310 bp band indicated the *Pcsk9* knockout allele and a 610 bp band indicated the wild-type allele. To genotype the *Cdh23^ahl^* allele, a second PCR was performed using 1 µL of genomic DNA and the primers 5′- CTAGAGAACCCACGCAGGAC-3′ and 5′-TCAGCCCAAGCTTCTACTGT-3′. The PCR protocol included an initial denaturation at 96°C for 2 minutes, followed by 30 cycles of denaturation at 96°C for 30 seconds, annealing at 52°C for 30 seconds, and extension at 68°C for 45 seconds. Two microliters of the resulting PCR product were digested with the restriction enzyme BsrI for 1 hour at 65°C, and the digestion products were separated on 2% E-Gel™ agarose gels. The presence of a 364 bp digestion product indicated successful replacement of the C57BL/6J-derived *Cdh23^ahl^* allele with the wild-type *Cdh23^+^* allele.

### Auditory Testing

Distortion product otoacoustic emissions (DPOAEs) and auditory brainstem responses (ABRs) were measured prior to cisplatin administration and following completion of the cisplatin protocol, as previously described (Fernandez et al. 2020; Fernandez et al. 2019). Briefly, mice were anesthetized using ketamine (Pulney Inc., Portland, ME, USA; 100 mg/kg IP) and xylazine (Akorn Inc., Lake Forest, IL, USA; 10 mg/kg IP). Core body temperature was maintained at 37°C using a temperature-controlled heating pad (World Precision Instruments ATC-2000, Sarasota, FL, USA). All auditory testing was conducted in a sound-attenuated booth (Acoustic Systems, Austin, TX, USA). DPOAE and ABR measurements were recorded using Tucker-Davis Technologies (TDT; Alachua, FL, USA) hardware (RZ6 Processor) and software (BioSigTZ).

For DPOAE testing, a single acoustic assembly consisting of an ER-10B+ microphone (Etymotic, Elk Grove Village, IL, USA) connected to two TDT MF-1 transducers was inserted into the external ear canal. An unobstructed path from port to tympanic membrane was established with an appropriate acoustic seal. Two primary tones were presented at 6 frequency pairs (f2 = 8, 11.2, 16, 22.4, 32, 40 kHz; f2/f1 = 1.2 from 30 to 90 dB SPL in 5 dB steps. DPOAE amplitudes at 2f1–f2 and background noise estimates were recorded.

ABR testing was conducted using tone-burst stimuli (Blackman window, 3 ms, 1.5 ms rise/fall) with alternating polarity at a presentation rate of 29.9/sec at 8, 11.2, 16, 22.4, 32, and 40 kHz. Signals were delivered via a closed-field TDT MF-1 speaker placed in the left ear of each mouse. Responses were recorded from subdermal needle electrodes (Rhythmlink, Columbia, SC, USA) placed at the vertex (noninverting), under the test ear (inverting), and at the base of the tail (ground). Waveforms from 1,024 presentations were acquired, averaged, amplified (20x), filtered (0.3-3 kHz) and digitized (25 kHz). Stimuli at each frequency were presented initially at 90 dB SPL, and decreased in 10 - 20 dB steps, and then in 5 dB steps near threshold. Threshold was determined by visual inspection and defined as the lowest stimulus level at which any identifiable and repeatable ABR wave was observed.

### Blood Collection

Blood samples were collected at designated time points before and after the cisplatin treatment protocol for subsequent analysis of plasma cholesterol levels. After briefly restraining the mice to facilitate venipuncture, the submandibular area was disinfected with alcohol, the submandibular vein located, punctured using a sterile 3–5 mm lancet. Approximately 150–250 μL of blood was collected into a 1.5 mL Eppendorf tube pre-loaded with 10–20 μL of heparin (Stemcell Technologies, cat# 07980) to prevent coagulation. Blood samples were placed on ice for 20 minutes prior to centrifugation at 2,000 × g for 10 minutes at 4°C. After centrifugation, 50–100 μL of plasma was carefully isolated and either used immediately for fluorometric total cholesterol assays or stored at −80°C for later analysis.

### Plasma Cholesterol Assays

Plasma cholesterol levels were quantified using the HDL and LDL/VLDL Cholesterol Assay Kit (Cat# STA-391, Cell Biolabs, Inc.) according to the manufacturer’s instructions. This enzymatic assay relies on a peroxidase-based reaction that generates a colorimetric signal, which was detected at 570 nm using a 96-well microtiter plate (Part# 234501, Cell Biolabs, Inc.) and measured with a fluorescence microplate reader (Vantastar, BMG Labtech).

A cholesterol standard curve was prepared using a series of eight cholesterol standard dilutions with final concentrations ranging from 0 to 12 µM per well. For measuring cholesterol levels in mouse plasma, 1 µL of plasma was diluted in 199 µL of 1X assay diluent buffer (Part# 239002, Cell Biolabs, Inc.) and aliquoted in triplicate into a 96-well plate, with a final volume of 50 µL per well. A 50 µL working solution was freshly prepared, consisting of 46.8 µL of 1X assay buffer, 1 µL of cholesterol oxidase (Part# 239004), 1 µL of HRP (Part# 234402), 1 µL of fluorescence probe (Part# 239005), and 0.2 µL of cholesterol esterase (Part# 239003). This solution was then added to each well, and the plate was incubated at 37°C for 45 minutes with gentle shaking while being protected from light.

Following incubation, fluorescence was measured using a fluorescence microplate reader (Vantastar, BMG Labtech) with an excitation wavelength of 530–570 nm and an emission detection range of 590–600 nm. Cholesterol concentrations were determined using the standard curve.

### Microdissection and Immunostaining

To estimate the total population of hair cells (HCs) in the cochlea after cisplatin treatment, whole-mount dissections of the organ of Corti were prepared from the right cochlea of each mouse. After intracardiac perfusion with 0.1M PBS, the cochleae were dissected from the temporal bone and intralabyrinthine perfused with 4% PFA. The following day, cochleae were incubated in 0.1M EDTA for 3–5 days at 4°C to allow decalcification, and the organ of Corti was microdissected under a dissecting microscope (Stemi 508, Zeiss) into 3–5 pieces. For each piece, the tectorial membrane was removed with forceps, and the spiral ligament was trimmed using an ophthalmic straight/Sideport Knife (Crest Point Ophthalmic).

Microdissected cochlear samples were collected and washed in 1× PBS, then preincubated in 30% sucrose for 30 minutes. To permeabilize the tissue, samples were placed on dry ice for 20 minutes and subsequently returned to room temperature for 30 minutes. After three washes in 1× PBS, samples were incubated for 1 hour at 37°C in blocking buffer consisting of 10% horse serum and 0.1% Triton X-100 in 0.1 M PBS. Primary antibody labeling was performed using anti-myosin VIIa (Proteus BioSciences, Inc., Ramona, CA; cat# 25-6790, lot no. 030118) diluted 1:200, and incubated at 37°C. The following day, samples were incubated with a donkey anti-rabbit Alexa Fluor 647 secondary antibody (Invitrogen, cat# A31573) at a 1:1000 dilution. After final washes in 1× PBS, samples were mounted on glass slides using Vectashield antifade mounting medium (Invitrogen) for imaging. Cochlear regions corresponding to defined frequency locations (ranging from 8 kHz at the apical turn to 40 kHz at the basal turn) were identified along the cochlear spiral using the Cochlear Frequency Mapping plug-in available from the Eaton-Peabody laboratory website. Images of each frequency region were acquired using an LSM900 laser scanning confocal microscope (Zeiss) with a 20× objective. Each image was digitally magnified ×1.7, reaching a final ×34 magnification. Hair cell counts were normalized to a standard-length segment (250 µm) to allow comparison across different cochlear regions.

### Statistics

All statistical analyses were performed in Prism 10.2.0 (GraphPad Software, Boston, MA), unless otherwise specified. Two-way ANOVA followed by Tukey’s multiple comparisons test was used to analyze both auditory threshold shifts (ABR and DPOAE) and hair cell counts across control and cisplatin-treated groups. Associations between baseline plasma cholesterol levels and auditory threshold shifts were assessed using simple linear regression. To further evaluate the effects of genotype and sex on high-frequency threshold shifts (32–40 kHz), Ordinary Least Squares (OLS) regression analysis was conducted in Python Data and are presented as mean ± SEM. Statistical significance is indicated as: (*) P < 0.05, (**) P < 0.01, (***) P < 0.001, (****) P < 0.0001. P-values above 0.05 (P > 0.05) were considered non-significant.

## 3. RESULTS

We previously reported that lovastatin conferred significant protection against CIHL in mice (Fernandez et al., 2020), and we used observational data to show that atorvastatin is associated with reduced CIHL among adults with head and neck cancer (Fernandez et al., 2021). To verify that our mouse model of CIHL (Fernandez et al., 2019) is consistent with our observational data in humans (Fernandez et al., 2021), we tested whether atorvastatin confers protection in mice. Cisplatin-treated CBA/CaJ mice underwent a three-cycle cisplatin administration protocol (Figure 1A). Atorvastatin-treated mice received once-daily atorvastatin via oral gavage beginning 24 hrs prior to cisplatin administration and then once daily throughout the cisplatin protocol. Cisplatin administration alone resulted in significant hearing loss (indicated by ABR threshold shifts) at all tested frequencies (Figure 1B). High-frequency (22.4 – 40 kHz) DPOAE threshold shifts were also evident in mice receiving cisplatin (Figure 1C), indicating significant outer hair cell (OHC) dysfunction. Concurrent administration of atorvastatin at 40 mg/kg or 60 mg/kg with cisplatin resulted in significant protection against cisplatin-induced ABR threshold shifts at 8 to 32 kHz (p ≤ 0.05, Figure 1B) and DPOAE threshold shifts at 8, 22.4, and 32 kHz (p ≤ 0.05, Figure 1C). These data indicate that atorvastatin significantly protects against CIHL and OHC dysfunction in mice, consistent with our observational data in humans (Fernandez et al., 2021).

The mechanisms underlying the protective effect of statins against CIHL are unknown; however, the hallmark activity of statins is inhibition of HMG-CoA reductase, which catalyzes the rate-limiting step in cholesterol synthesis. This inhibition of HMG-CoA reductase by statins results in reduced plasma cholesterol levels. We hypothesized that reduced plasma cholesterol could offer protection against hearing loss, even in the absence of a statin. To test this hypothesis, we utilized *Pcsk9* knockout (KO) mice (B6;129S6-Pcsk9^tm1Jdh^/J) as a genetic model with reduced plasma cholesterol.

*Pcsk9 KO* mice exhibit plasma cholesterol levels approximately 50% lower than their wild-type (WT) littermates (Rashid et al., 2005). However, these data were obtained in mice maintained on a C57BL/6 (B6) background, which carry a mutation in the gene encoding cadherin 23, *Cdh23^ahl^*, that causes hearing loss beginning as early as 12 weeks of age (Johnson et al., 2017). To determine whether *Pcsk9* deletion continues to result in reduced cholesterol levels after correction of the *Cdh23^ahl^* allele that results in premature hearing loss, we assessed total plasma cholesterol levels at baseline and following cisplatin exposure in both male and female B6.CAST-Pcsk9^tm1Jdh^ Cdh23^Ahl+^ mice hereafter referred to as *Pcsk9 KO* mice and WT littermates. Prior to initation of cisplatin administration, plasma was collected and total cholesterol was measured using a colorimetric assay (Figure 2A). *Pcsk9 KO* mice exhibited significantly reduced serum cholesterol levels compared to WT littermates. No significant sex differences in cholesterol levels were observed within either genotype as determined by two-way ANOVA. To assess whether cisplatin exposure affects systemic cholesterol levels, plasma was collected after completion of the cisplatin administration protocol and total cholesterol was measured. The relative difference in plasma cholesterol levels between *Pcsk9 KO* and WT mice remained (Figure 2B) and no sex-specific effects were detected post-treatment (Table 1). These results confirm that *Pcsk9* deletion leads to reduction in plasma cholesterol levels, independent of sex or cisplatin exposure. Therefore, *Pcsk9 KO* mice are a suitable genetic model to test whether reduced cholesterol protects against CIHL.

**Figure 2.**
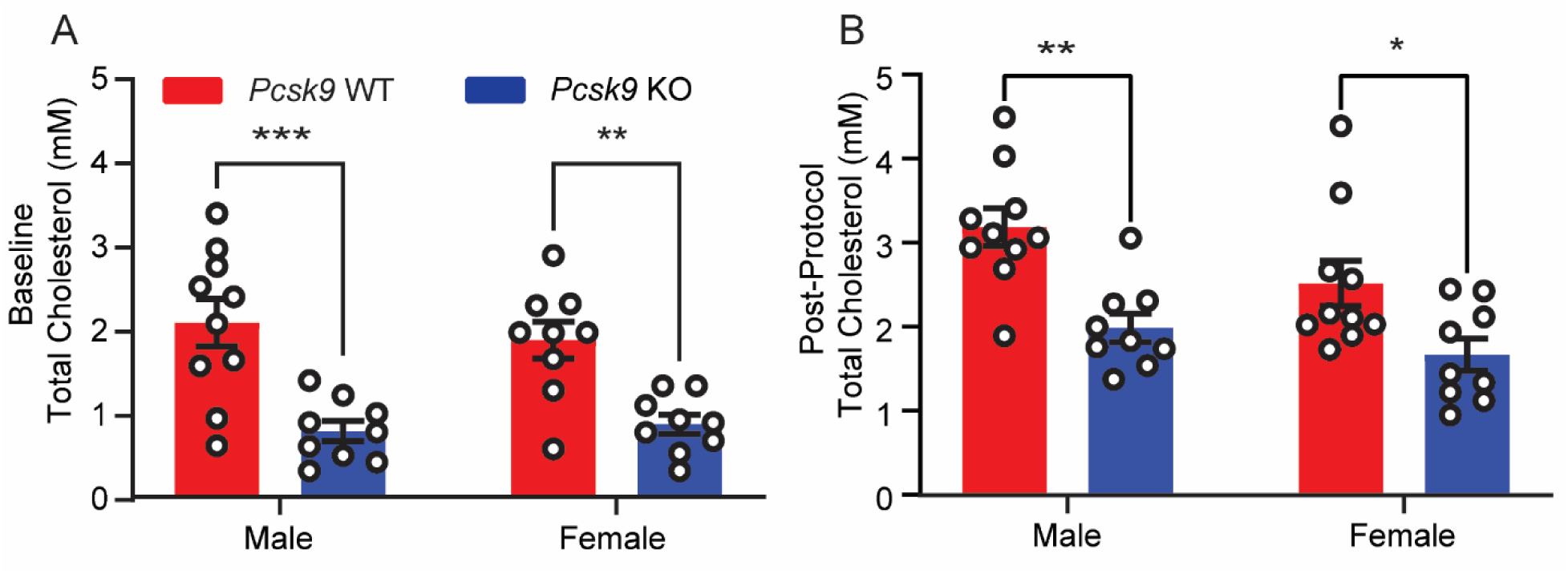
*Pcsk9 KO* mice have reduced total cholesterol levels compared to WT mice. **(A)** Total plasma cholesterol measured at baseline in male (n = 19) and female (n = 18) mice, organized by genotype. **(B)** Total plasma cholesterol measured post-cisplatin treatment in male (n = 19) and female (n = 19) mice, organized by genotype. At both timepoints, *Pcsk9 KO* mice had significantly lower cholesterol levels compared to WT mice as determined by two-way ANOVA, [* p ≤ 0.05, ** p ≤ 0.01, *** p ≤ 0.001].

**Table 1:**
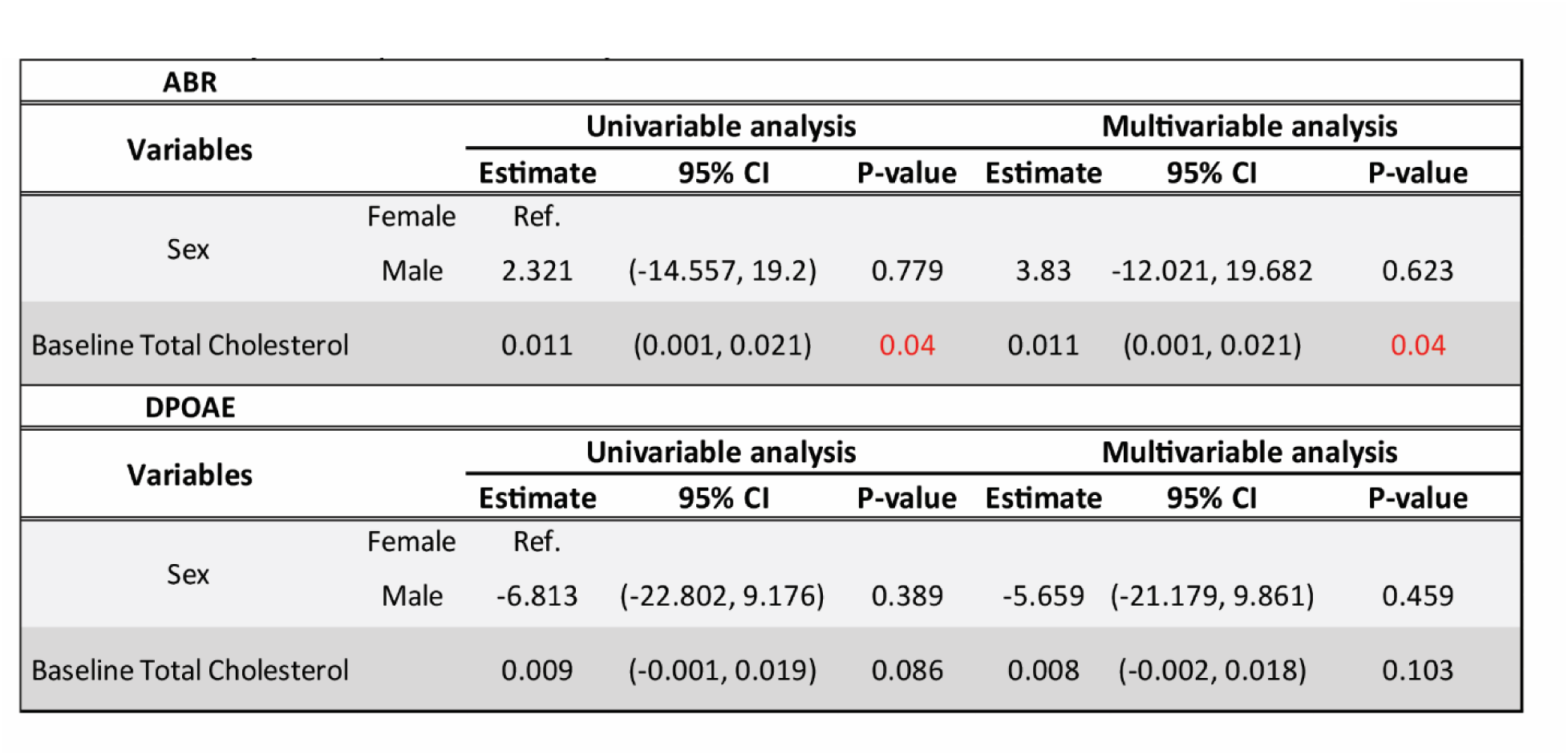
Ordinary Least Squares (OLS) analysis for ABR and DPOAE threshold shift at 32-40 kHz.

To determine if *Pcsk9 KO* mice are protected against CIHL, auditory function was assessed by measuring ABR and DPOAE thresholds before and after three cycles (Figure 1A) of cisplatin treatment. Saline-treated WT and *Pcsk9 KO* mice exhibited no hearing loss, as measured by ABR threshold shifts (Figure 3A). WT mice treated with cisplatin exhibited significant cisplatin-induced ABR threshold shifts at 32 and 40kHz, consistent with the characteristic high-frequency hearing loss observed clinically (Fernandez et al., 2021; Frisina et al., 2016; Marnitz et al., 2018). In contrast, *Pcsk9 KO* mice showed significantly less cisplatin-induced hearing loss compared to WT mice (Figure 3A). Similarly, saline-treated WT or *Pcsk9 KO* mice and cisplatin-treated *Pcsk9 KO* mice showed no significant difference in outer hair cell function as measured by DPOAE threshold shifts (Figure 3B). In contrast, WT mice treated with cisplatin exhibited significant DPOAE threshold shifts at 32 and 40kHz (Figure 3B). Together these data indicate that compared to WT mice, *Pcsk9 KO* mice are protected against CIHL (as measured by ABR threshold shifts) and OHC dysfunction (as measured by DPOAE).

**Figure 3.**
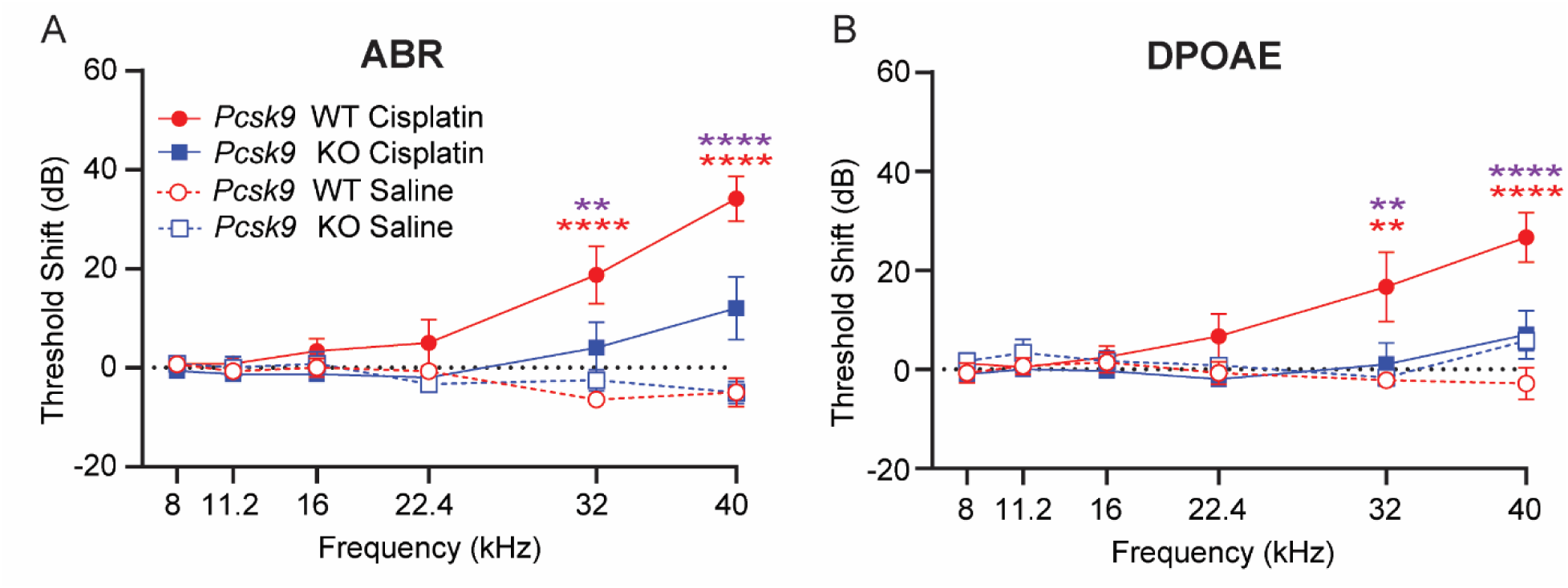
*Pcsk9 KO* mice are resistant to CIHL and OHC dysfunction. **(A)** ABR threshold shifts were measured in saline-treated WT (n = 7) and KO (n = 6) mice, as well as in cisplatin-treated WT (n = 12) and KO (n = 15) mice. Cisplatin-treated WT mice had significant CIHL (ABR threshold shifts) at 32 and 40 kHz (****p < 0.0001, two-way ANOVA) compared to saline-treated mice. In contrast, cisplatin-treated *Pcsk9 KO* mice had reduced CIHL compared to WT mice at 32 kHz (**p = 0.0055, two-way ANOVA) and 40 kHz (****p < 0.0001, two-way ANOVA). **(B)** Consistent with our observations of reduced ABR threshold shifts, cisplatin-treated *Pcsk9 KO* mice had reduced DPOAE threshold shifts in comparison to cisplatin-treated WT mice at 32 kHz (**\*\***p = 0.0015) and 40 kHz (**\*\*\*\***p < 0.0001) in post-hoc comparisons following a two-way ANOVA. No sex differences in CIHL severity were observed, and saline-treated control mice of both genotypes exhibited no significant changes in ABR or DPOAE thresholds. Data are expressed as mean ± SEM [Purple asterisks: Cisplatin-treated *Pcsk9 KO* vs. Cisplatin-treated WT; red asterisks: Cisplatin-treated WT vs. saline-treated WT, as determined by two-way ANOVA, ** p ≤ 0.01, **** p ≤ 0.0001].

We next asked whether the severity of CIHL was correlated with plasma cholesterol levels. To explore this, we compared the mean high-frequency threshold shift (32 and 40 kHz)—measured by both ABR and DPOAE—with total plasma cholesterol levels at baseline. In cisplatin-treated mice, high-frequency ABR threshold shifts showed a significant correlation with baseline cholesterol levels (p = 0.04; Figure 4A, Table 1). DPOAE threshold shifts at the same frequencies did not show a significant correlation (p = 0.086; Figure 4B, Table 1). Univariable and multivariable regression analyses (Table 1) reinforced the significant association between baseline cholesterol levels and high-frequency ABR threshold shifts, independent of sex. These results are consistent with a model in which reduced plasma cholesterol confers protection against CIHL.

**Figure 4.**
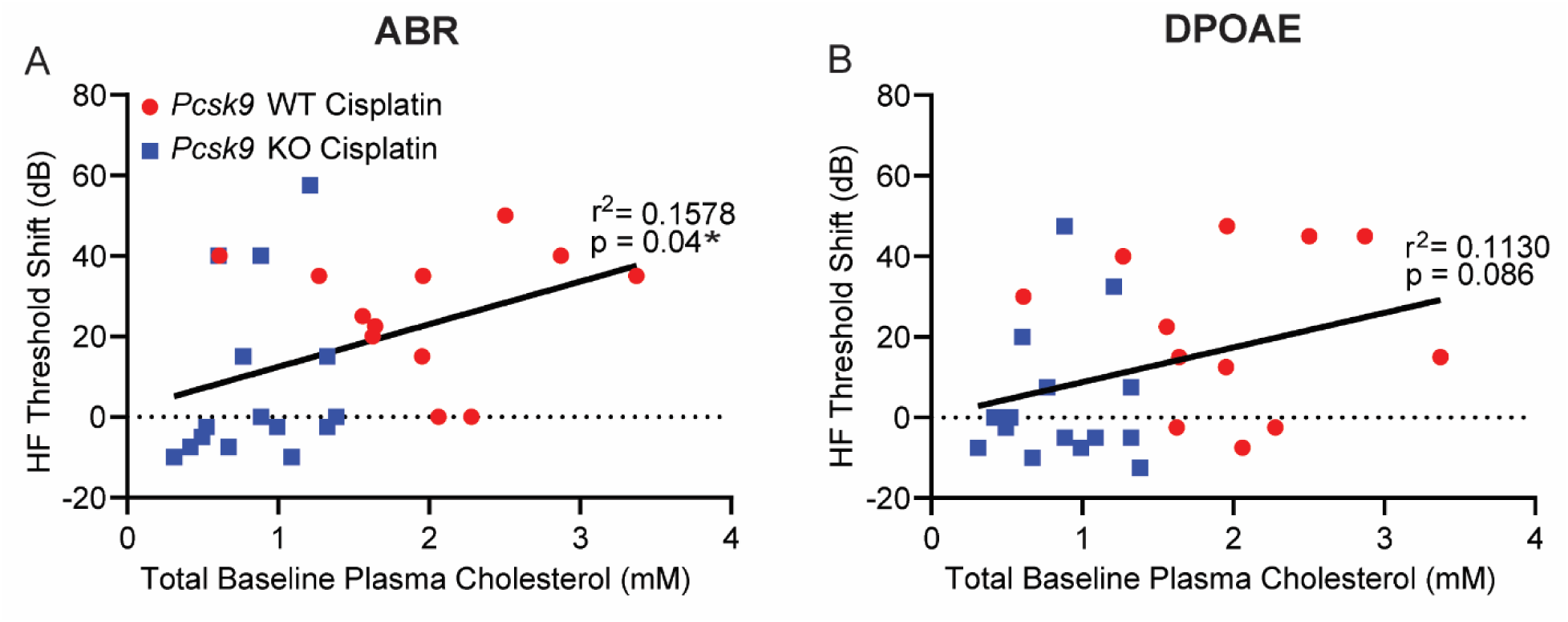
The severity of CIHL is correlated with plasma cholesterol levels. High-frequency threshold shifts (average of 32 and 40 kHz), measured by ABR and DPOAE, were analyzed in relation to baseline plasma cholesterol levels in cisplatin-treated WT (n = 12) and *Pcsk9 KO* (n = 15) mice using simple linear regression. ABR threshold shifts showed a significant correlation with baseline cholesterol levels (*p = 0.04; Figure 4A), whereas DPOAE threshold shifts did not significantly correlate with plasma cholesterol levels (p = 0.086; Figure 4B).

Following completion of auditory testing, cochleae from *Pcsk9 KO* and WT mice treated with saline or cisplatin were harvested, micro dissected, and labeled with an antibody against MYO7A, a hair cell marker (Figure 5A). Saline-treated cochleae showed three rows of intact outer hair cells (OHCs) and a single row of intact inner hair cells (IHCs). Cisplatin resulted in observable loss of OHCs in WT control mice, especially at the most vulnerable cochlear regions corresponding to higher frequencies (Figure 5A, 5B). In contrast, cisplatin-treated *Pcsk9 KO* mice were protected against cisplatin-induced OHC death (Figure 5A, 5B). No differences in IHC survival were observed. OHC loss was consistent with the observed ABR and DPOAE threshold shifts.

**Figure 5.**
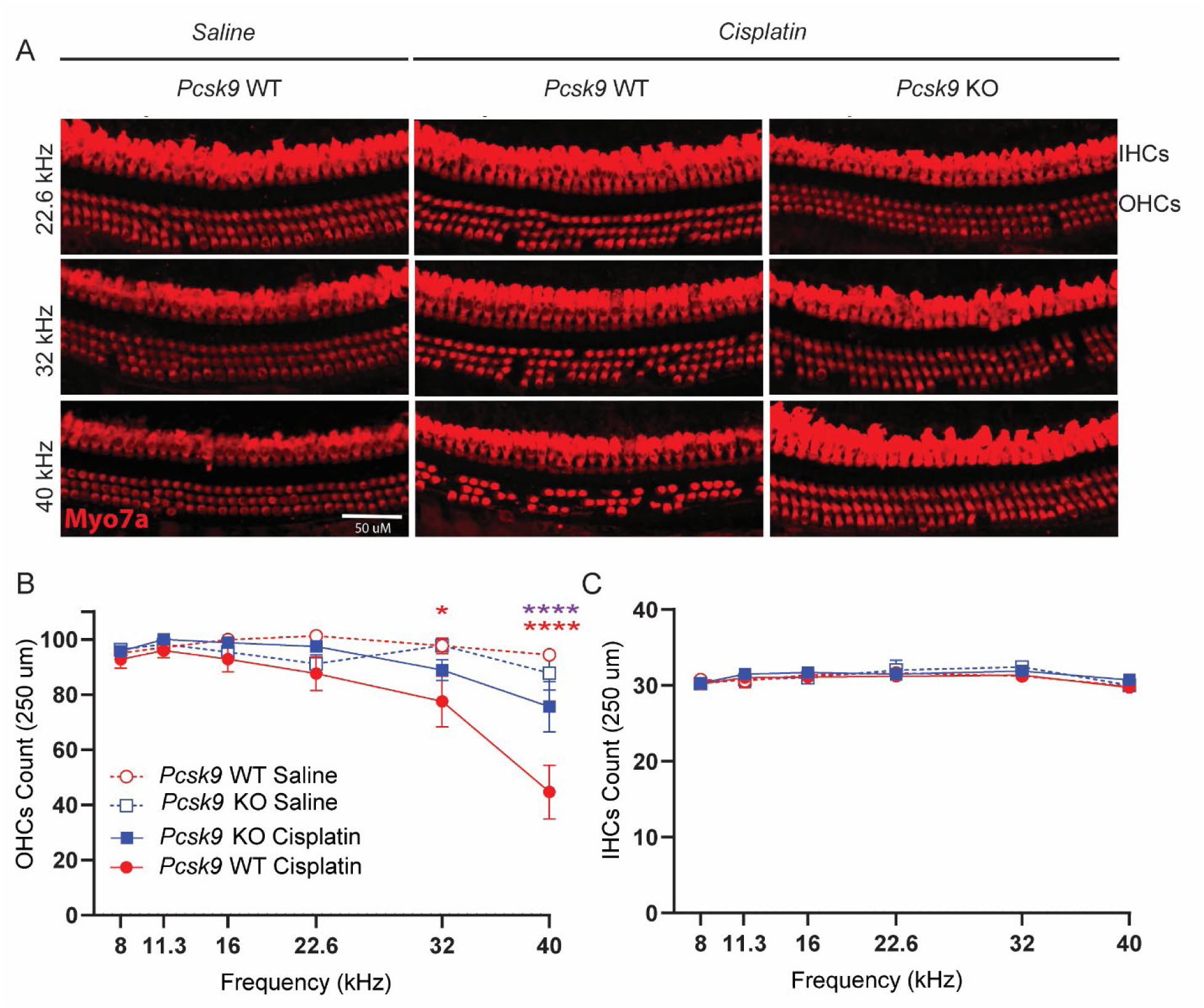
*Pcsk9 KO* mice are protected against cisplatin-induced outer hair cell (OHC) death. **(A)** IHCs and OHCs from *Pcsk9 KO* and WT mice treated with saline or cisplatin were labeled with an antibody against *myosin7a* (MYO7A). Shown are representative images from three cochlear regions, corresponding to 22.6 kHz (top row), 32 kHz (middle row), and 40 kHz (bottom row). IHCs and OHCs from WT mice treated with cisplatin (n = 11), only show OHC loss at 22.6 kHz, 32 kHz and 40 kHz regions. Cisplatin-induced OHC loss at these frequency regions was reduced in *Pcsk9 KO* mice (n = 15). **(B)** Cochleae from WT (n = 8) and *KO* (n = 5) mice treated with saline showed no OHC loss. Cisplatin resulted in significant OHC death in WT mice at 32 kHz (*p = 0.0362, two-way ANOVA) and 40 kHz (****p < 0.0001, two-way ANOVA). Cisplatin-treated *Pcsk9 KO* mice were significantly protected against cisplatin-induced OHC death at 40kHz (****p < 0.0001, two-way ANOVA). **(C)** No differences in IHC survival were observed among any of the groups. Data are expressed as mean ± SEM [Purple asterisks: Cisplatin-treated *Pcsk9 KO* vs. Cisplatin-treated WT, red asterisks: Cisplatin-treated WT vs. saline-treated WT; as determined by two-way ANOVA, * p ≤ 0.05, **** p ≤ 0.0001].

## 4. DISCUSSION

Cisplatin is an essential chemotherapeutic agent due to its efficacy against a broad spectrum of malignancies; however, it results in permanent hearing loss in over half of treated adults. There is a critical unmet clinical need for therapies to reduce CIHL in adults. Our data in mice (Figure 1) and observational data in humans (Fernandez et al., 2021) suggest that atorvastatin, an FDA-approved drug with a good safety profile, holds potential to be repurposed to reduce the incidence and severity of CIHL. The mechanisms underlying the protective effect of atorvastatin remain unclear; however, the hallmark activity of atorvastatin is inhibition of HMG-CoA reductase, resulting in reduced plasma cholesterol levels. Here we used *Pcsk9 KO* mice as a genetic model of reduced plasma cholesterol to test the hypothesis that reduced plasma cholesterol is protective against CIHL independent of statin treatment. We find that *Pcsk9 KO* mice have reduced plasma cholesterol compared to WT littermates, and that *Pcsk9 KO* mice are resistant to CIHL and OHC dysfunction and death. Together these data indicate that reduction of plasma cholesterol is protective against CIHL, and they are consistent with a model in which the protective effect of statins against CIHL is mediated by reduced serum cholesterol.

The protective phenotype observed in *Pcsk9 KO* mice suggests a cholesterol-dependent mechanism of cochlear vulnerability to cisplatin that impacts OHC endurance. Notably, we found a statistically significant correlation between high-frequency ABR threshold shifts and total cholesterol levels, consistent with the idea that cholesterol regulation influences susceptibility to CIHL.

These findings raise an important question regarding the underlying mechanisms of protection against CIHL observed in *Pcsk9 KO* mice. While our data cannot rule out the possibility that absence of PCSK9 contributes directly to cochlear protection, the simplest interpretation of our data is that this protection is mediated through reduced systemic cholesterol levels—a well-established consequence of PCSK9 inhibition (Bao et al., 2024; Bardolia et al., 2021; Chen & Chen, 2023; Sabatine et al., 2017; Sobati et al., 2020). Supporting this interpretation, we also observed that atorvastatin, a widely used cholesterol-lowering drug, similarly protects against CIHL in mice.

Our data do not address whether the protective effect observed in *Pcsk9 KO* mice is mediated locally (in the inner ear) or systemically. PCSK9 is a circulating protein best known for its role in hepatic regulation of LDLR, and its function within the cochlea is unknown. The cochlea is protected by the blood-labyrinth barrier (BLB), which restricts the entry of large circulating molecules into the endolymphatic compartment.

This is particularly relevant for the scala media, which houses the organ of Corti and the stria vascularis. Due to the selective permeability of the BLB, it is unlikely that circulating PCSK9 can reach these compartments in significant amounts. While PCSK9 is present in cerebrospinal fluid (Chen et al., 2014; Courtemanche et al., 2018) and could potentially access the scala tympani through the perilymph, its ability to penetrate the scala media appears limited. If PCSK9 is indeed regulating LDLR in regions such as the stria vascularis or organ of Corti, it raises the possibility that the protein might be synthesized locally within the cochlea. Transcriptomic datasets available on the Gene Expression Analysis Resource (gEAR) resource (Orvis et al., 2021) and our own unpublished single nucleus RNA-seq data suggest that *Pcsk9* mRNA may be expressed in cochlear hair cells. Clarifying whether PCSK9 acts locally or systemically will be essential for understanding its role in cochlear lipid regulation and its potential in cochlear protection.

LDL receptors are expressed broadly within the cochlea (Saume et al., 2021), where they may mediate the uptake of cholesterol into several cell types. PCSK9 facilitates the degradation of LDL receptors, and thus, genetic deletion of PCSK9 prolongs the presence of LDL receptors at the cell surface, enhancing LDL-C clearance from the circulation and resulting in reduced plasma LDL-C levels. The cellular uptake of cholesterol, mediated by LDLR, may modulate membrane fluidity and stiffness (Saume et al., 2021). Functionally, cholesterol dynamics within the cochlea influence ion channel activity and membrane properties as well as hair cell mechanotransduction and electromotility, processes that are essential for generating and maintaining the endocochlear potential, the electrochemical driving force for hair cell mechanotransduction (Brownell et al., 2011; Purcell et al., 2011; Saume et al., 2021). As demonstrated in this and previous studies, cisplatin induces significant OHC death, strial dysfunction, and diminished endocochlear potential (Breglio et al., 2017). Given that hepatocytic LDLR expression is increased in *Pcsk9 KO* mice (Roubtsova et al., 2022), it is possible that LDLR levels in the stria vascularis, spiral ligament, and supporting cells of the organ of Corti are also increased. These increases may result in reduced cellular cholesterol levels, which could alter the membrane properties (Alobeedallah et al., 2020) and the activity of membrane-regulated proteins such us the ion channels responsible of the generation of the endocochlear potential (Brownell et al., 2011), mitigating the vulnerability of the inner ear to cisplatin-induced damage.

Our data support a model in which cholesterol homeostasis—regulated either pharmacologically through statins or genetically via PCSK9 deficiency—critically influences cochlear susceptibility to cisplatin. This study also highlights lipid-lowering strategies as a viable approach for CIHL prevention and encourages further investigation into the repurposing of cholesterol modulators for auditory protection.

## 5. CONCLUSION

Our study demonstrates that reduction of serum cholesterol through either pharmacologic or genetic means is protective against CIHL in mice. These data indicate that systemic cholesterol reduction increases resistance to cisplatin-induced hearing loss, and they suggest that the observed protective effect of atorvastatin against cisplatin-induced damage is mediated by cholesterol modulation.

## 6. ACKNOWLEDGMENTS

We thank Dr. Angela Ballesteros and Dr. Meghan Drummond for their helpful comments on and review of the manuscript. We also thank Dr. Drummond for valuable insights on the mouse model utilized in this project and Sherly Michel (NIDCD) for her assistance with submandibular blood draws. We acknowledge the support of the Mouse Auditory Testing Core at the National Institute on Deafness and Other Communication Disorders (NIDCD), National Institutes of Health, which is funded through project number ZIC DC000080.

## 7. DISCLAIMER

This research was supported [in part] by the Intramural Research Program of the National Institutes of Health (NIH). The contributions of the NIH author(s) are considered Works of the United States Government. The findings and conclusions presented in this paper are those of the author(s) and do not necessarily reflect the views of the NIH or the U.S. Department of Health and Human Services.

---

**Supplementary Figure 1.**
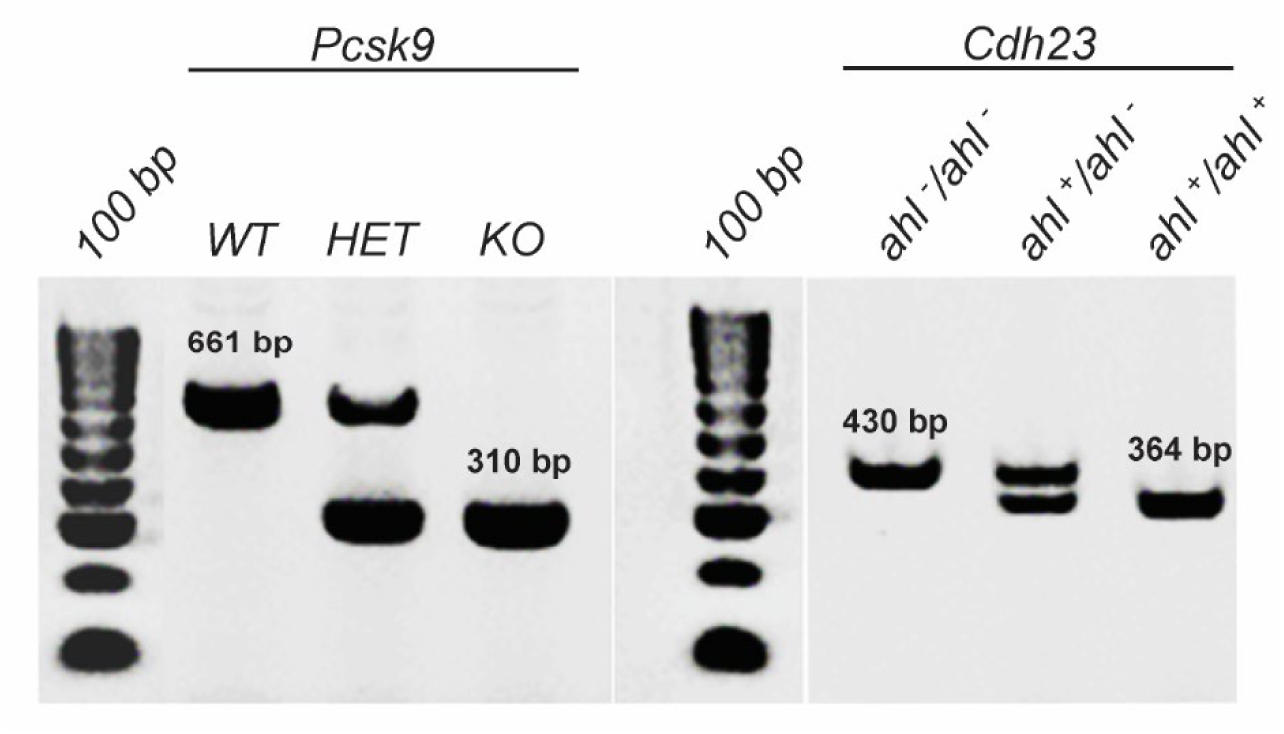
PCR-based genotyping followed by BsrI restriction digestion. A 310 bp band confirmed the deletion of *Pcsk9*, while the presence of a BsrI digestion product at 364 bp indicated successful replacement of the *Cdh23^ahl^* locus with the wild-type *cast*-derived *Cdh23* allele.

