## Supplementary Figure 1 for "Effects of Cholesterol Modulation on Cisplatin-Induced Hearing Loss"

Regeneron - Internal

**Supplementary Figure 1**

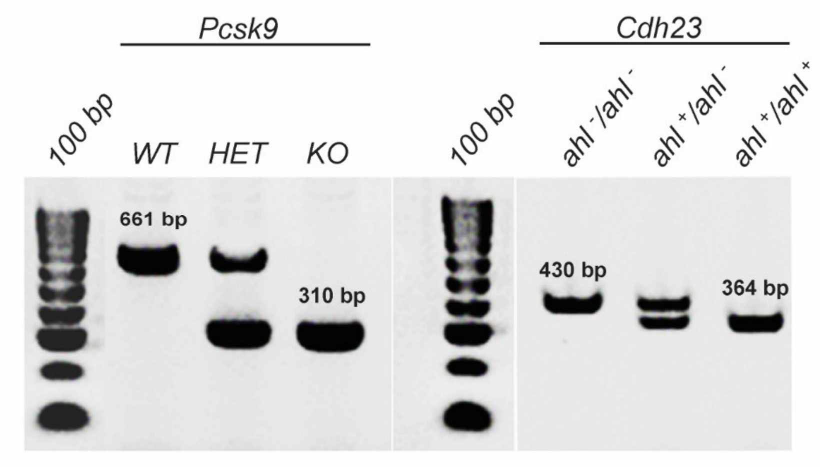


**Figure 1** – PCR-based genotyping followed by Bsr**I** restriction digestion. A 310 bp band confirmed the deletion of *Pcsk9*, while the presence of a Bsr**I** digestion product at 364 bp indicated successful replacement of the *Cdh23*$^{ahl}$ locus with the wild-type *cast*-derived *Cdh23* allele.
